# Establishing an AI-based evaluation system that quantifies social/pathophysiological behaviors of common marmosets

**DOI:** 10.1101/2023.10.16.561623

**Authors:** Takaaki Kaneko, Jumpei Matsumoto, Wanyi Lu, Xincheng Zhao, Louie Richard Ueno-Nigh, Takao Oishi, Kei Kimura, Yukiko Otsuka, Andi Zheng, Kensuke Ikenaka, Kousuke Baba, Hideki Mochizuki, Hisao Nishijo, Ken-ichi Inoue, Masahiko Takada

**Affiliations:** Center for the Evolutionary Origins of Human Behavior, Kyoto University, Inuyama, Aichi 484-8506, Japan; Department of System Emotional Science, Faculty of Medicine, University of Toyama, Toyama 930-0194, Japan; Research Center for Idling Brain Science, University of Toyama, Toyama 930-0194, Japan; Department of Neurology, Osaka University School of Medicine, Suita, Osaka 565-0871, Japan

## Abstract

Nonhuman primates (NHPs) are indispensable animal models by virtue of the continuity of behavioral repertoires across primates, including humans. However, behavioral assessment at the laboratory level has so far been limited. By applying multiple deep neural networks trained with large-scale datasets, we established an evaluation system that could reconstruct and estimate three-dimensional (3D) poses of common marmosets, a small NHP that is suitable for analyzing complex natural behaviors in laboratory setups. We further developed downstream analytic methodologies to quantify a variety of behavioral parameters beyond simple motion kinematics, such as social interactions and the internal state behind actions, obtained solely from 3D pose data. Moreover, a fully unsupervised approach enabled us to detect progressively-appearing symptomatic behaviors over a year in a Parkinson’s disease model. The high-throughput and versatile nature of our analytic pipeline will open a new avenue for neuroscience research dealing with big-data analyses of social/pathophysiological behaviors in NHPs.

## Introduction

Quantitative evaluation of animal behavior is crucial for various research areas of neuroscience. However, observing natural behaviors of freely moving animals by visual inspection incurs a considerable cost. Meanwhile, recent advances in artificial intelligence (AI) allow us to pave the way to quantify massive amounts of behavioral data in a large-scale and automated manner^1–4^, and assessment of natural behaviors with “markerless pose estimation” has already been implemented in a number of studies^5–10^. Indeed, AI-based three-dimensional (3D) analysis of body posture, involving the limb positions, makes it possible to evaluate a variety of behavioral aspects that characterize nonhuman primates (NHPs)^11,12^.

The application of this methodological innovation to the neuroscience research field is now rapidly expanding^11,13–17^, as it is expected to have a potential to bring about fundamental changes in how to design behavioral experiments on NHPs which have long been carried out in a head-fixed condition. In the past decades, accumulated evidence from a number of research works, such as ethological studies on wild animals, suggests the continuity of behavioral repertoires across primates including humans^19–21^. However, there remains a large gap between the field and the laboratory research since experimental settings under freely moving conditions have so far been limited at the laboratory level.

Common marmosets are one of the NHP species suitable for overcoming this problem, given that their relatively small body size permits observations of complex natural behaviors in laboratory setups^18–21^. Furthermore, marmosets are a remarkably prosocial animal. It is generally accepted that all family members cooperate to breed infants whose development is successfully attained via interactions with their caregivers. This implies that marmosets can be useful as a primate model for exploring social behavior^18,22^. The development of telemetric devices for brain activity recordings^28–30^ also accelerates the preparation of experimental environment in a freely behaving fashion. In addition, the utility of marmosets which have high reproductive efficiency has led to the production of brain disease models by genetic engineering techniques^23–27^, which requires the longitudinal and high-throughput assessment of symptomatic behaviors.

Two issues should be solved to achieve a methodological improvement in designing behavioral experiments on marmosets. First, the practical use of “deep neural networks” for behavioral analysis demands both a huge volume of ground truth data ^14,16^ and an analytic pipeline that reconstructs 3D poses of multiple animals simultaneously while recognizing individuals. Second, even if the best effort is made to establish such a system, a major question still arises as to how effective this approach is to evaluate natural behaviors of freely moving marmosets. In fact, quantitative analyses to date based on the markerless pose estimation have highly been focused on the movement itself (e.g., kinematics of body-part movements and sequence of motor actions) ^6,17,31^, leaving cognitive behaviors or social interactions untargeted.

In the present study, we developed a markerless 3D pose estimation system to analyze natural behaviors of marmosets under freely moving conditions, and a large-scale training dataset to promote automated quantification of videographic data. We further developed a set of downstream analytic methodologies that took advantage of the potential of 3D pose data. Here we show that (1) the 3D pose data are suitable for defining social behavior which should be more than kinematics of a single animal and represents complex interactions among multiple animals, (2) the 3D pose data are able to infer the animal’s internal state behind actions, and (3) a completely unsupervised approach based on the 3D pose data allows us to detect behavioral changes in response to pathophysiological conditions. Through these distinct experimental subjects (parenting behavior of male vs. female marmosets, behavioral flexibility of socially interacting marmosets, and symptomatic behaviors progressively appearing in a marmoset model of Parkinson’s disease (PD), respectively), we have revealed the potent applicability of our system that permits extracting a wide range of behavioral parameters beyond spatiotemporal kinematics.

## Results

### Markerless 3D pose estimation of multiple marmosets with individual identification

Our analytic framework consisted of the following three elements: a multi-camera recording system, an analytic pipeline combined with multiple deep neural networks, and large-scale ground truth data to train the deep neural networks for accurate quantification. The recording system included eight synchronized cameras surrounding a transparent cage that was specially designed to allow housing of a marmoset family (up to four individuals) and to provide continuous clear video recordings for several days or more. Multiview videographic data were fed into the custom-made analytic pipeline which had fully been optimized for robust reconstruction of the 3D poses of multiple marmosets under individual identification in a variety of natural behavioral contexts (Fig. 1a).

**Fig. 1.**
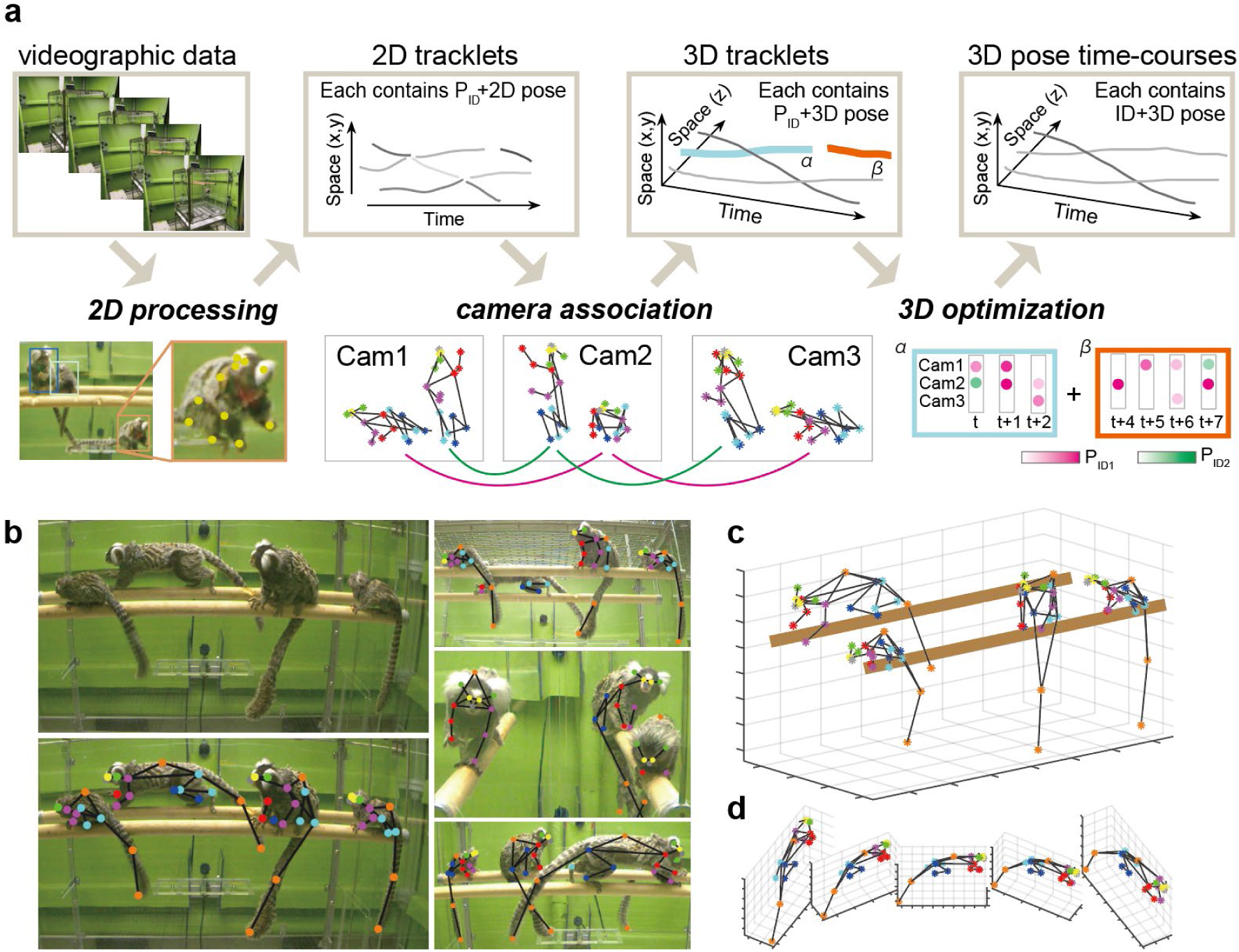
Analytic pipeline for AI-based quantification of marmoset natural behaviors. **a:** Overview of the analytic pipeline. For each of the videographic images captured by different cameras, short fragments of time-series data including postures and potential animal IDs, i.e., 2D tracklets, were generated (*2D processing*). Then, cross-view matching across the cameras was carried out to construct 3D tracklets representing 3D postures and potential animal IDs (*camera association*). Lastly, 3D pose time-courses for each animal were obtained by combining multiple 3D tracklets over the entire recoding time based on spatial continuity and animals IDs of the 3D tracklets (*3D optimization*). Note that the two spatial dimensions (x, y) are shown in one axis instead of two different axes, but this is only for visualization purposes. **b:** 3D annotations of a marmoset family. Upper-left, a cropped image taken from a single camera. Lower-left, the same image with the annotations. Right, cropped images of the same scene taken from different cameras. **c:** Reconstructed 3D poses of a marmoset family obtained from the same scene as b. **d:** Exemplified 3D poses from different viewpoints.

For the analytic pipeline, regions of interest (ROIs) where marmosets were located were first determined in each camera view at each time frame by using a detection network. Subsequently,18 keypoints and a potential animal identity per ROI were estimated through a pose network and an identity network, respectively. In each camera view, ROIs taken from numbers of time frames were combined based on the spatial continuity to construct tracklets which were composed of time-series data including the pose and identity. During this process, individual tracklets contained information only from a single camera view, and, therefore, they were fragmented by a short time period (Fig.1a, *2D processing*). As the next step, a 3D tracklet was constructed by combining several tracklets that represented the same animal from different camera views by minimizing the so-called pose affinity score (Fig. 1a, *camera association*; for details, see the Methods section). Finally, 3D tracklets were combined across the entire recording time based on both the spatial continuity and the probability of animal identity (Fig. 1a, *3D optimization*).

To achieve the accurate and robust 3D pose estimation, we created annotations of 3D keypoints for more than 7404 bodies (consisting of eight different views) (Fig. 1c, d) in a variety of natural behavioral contexts (Fig. 2a, b), which could be used as a ground truth dataset for training both the detection and pose networks. The requirement for a training dataset of animal recognition largely depended on experimental conditions (with/without infants, the use of a color tag, implantation of neuron activity recording devices, etc.). In the present study, we tested either a pair of marmosets or a breeding family (including male and female parents with their infants). A neckless type of color tag was attached to adult marmosets to facilitate identification. Under these conditions, we labeled 4231 samples in total for ID classification. With this dataset, we used 80% for training and 10% each for validation and test. The ground truth dataset was created from 29 different individuals ranging from 1.5 months old (infant) to 12 years old (adult).

**Fig. 2.**
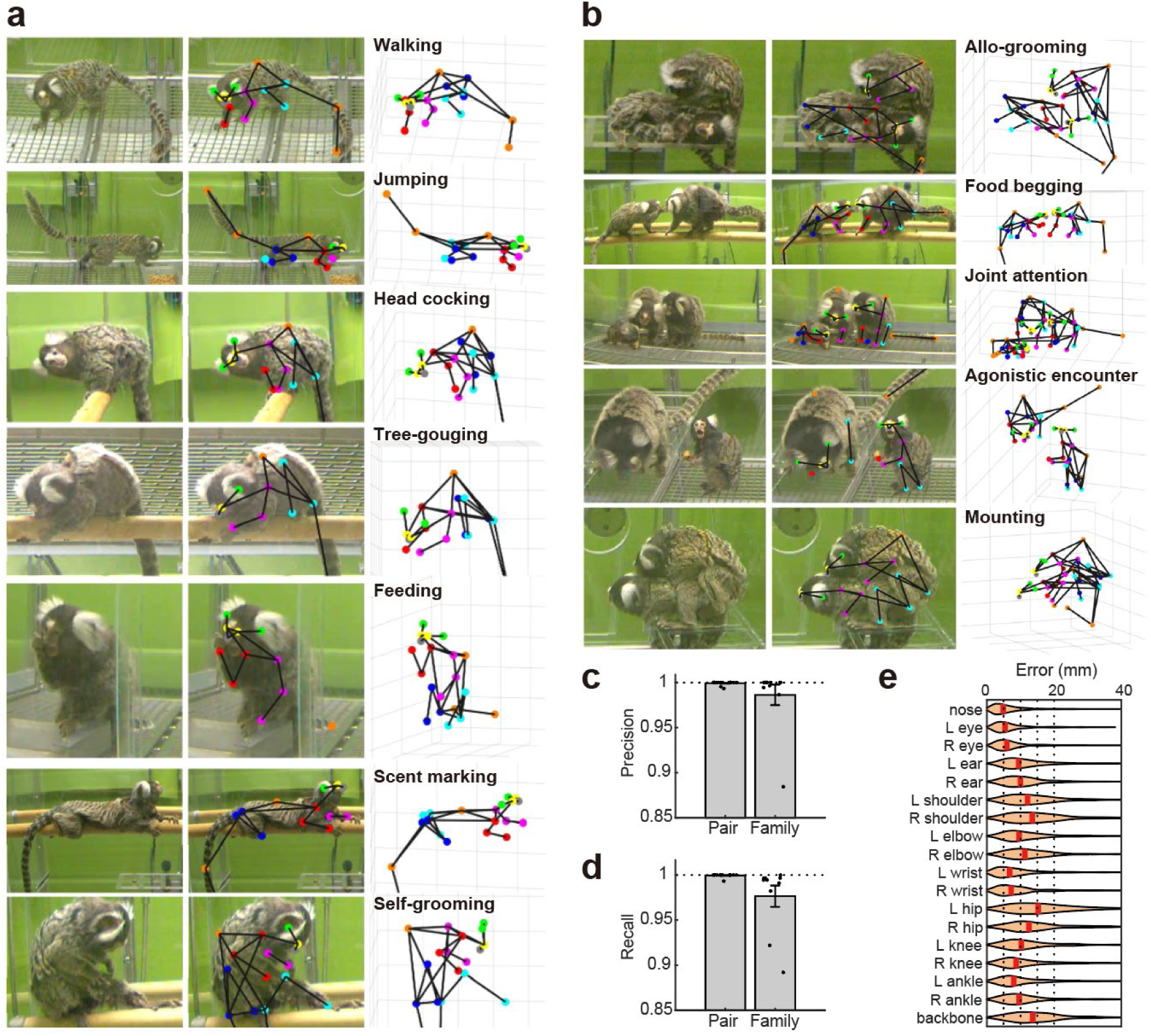
Exemplified annotations and decoding accuracy of the analytic pipeline. **a:** Examples of annotations in different behavioral contexts. Each column denotes the original images, the same images with annotations, and the 3D poses, respectively, from the left to the right. **b:** Examples of social behaviors. We obtained annotations for 2733 scenes, and 7404 and 56103 bodies in a 3D or 2D space, respectively. **c,d:** Precision and recall rates for detection and identification of individual animals. **e:** Errors between annotations and AI prediction of 3D poses. Errors were around 10 mm in marmosets with 200 mm of body length.

The final performance of animal detection and identification in 3D space was 99.3% and 98.8% in precision and recall, respectively (Fig. 2c, d, Video S1). The geometric error in pose estimation at each keypoint was 9.68 mm (4.86 ∼15.25) in 3D space (Fig. 2e). On the scale of human body, the estimation error of, for example, the wrist positions were about 4 cm. This accuracy was comparable to the state-of-the-art performance of a similar task in human pose estimation^32^ where enormous amounts of ground truth data were available, indicating that our system consisting of the recording environment, training dataset, and analytic pipeline reached the highest level that was considered achievable at the present time. However, a major question remained as to the extent to which our system would practically be useful for actual experiments, which was hard to judge from the so-far-listed score alone. In the following sections, we explored the potential of 3D pose data by quantifying various types of behavioral parameters that were beyond simple spatiotemporal kinematics of the body parts.

### Differential roles of male vs. female marmosets in parenting as defined by automated detection of social behavior

When introducing the automated quantification into natural behaviors, evaluation of social behavior is the most difficult and beneficial, since it is more than kinematics of a single animal. In the first set of our experiments, we tested the potential of 3D pose data for assessing food-sharing behavior which is frequently observed in a breeding family of marmosets. Both male and female marmosets generally take care of their infants together, and, therefore, they are characterized as cooperative breeders, which is similar to the human case, but is relatively rare in other NHPs^18^. As part of parenting, adult marmosets share their food with infant marmosets, which enables the infants not only to satisfy their nutritional needs, but also to obtain an opportunity of learning about diet^33^. Thus, we attempted to quantify food-sharing behavior of breeding marmoset families.

In the present experiment, we sought to detect the food-sharing behavior by applying a spatiotemporal filter to 3D pose time-course data. Two marmoset families participated in this experiment. Since the output of our system was a simple time-course data of the 3D posture in each marmoset, we started with engineering the features that might capture food-sharing events in the marmoset families based on the 3D pose time-course data. According to such data obtained from parents and infants, we computed the distance between the either the infants’ mouths/hands and those of parents’ hands/mouths, and its derivatives (i.e., velocity). Comparison with videographic images confirmed that the resulting time-course data could be potentially good indicators to detect the food-sharing event between the parents and the infants (Fig. 3a, b). Via spatiotemporal thresholds of these quantitative posture and motion parameters, we then defined and counted the occurrence of such events automatically (for details, see the Methods section). Moreover, we acquired annotations by a human observer to optimize and verify the automated detection of food-sharing events based on a subset of videographic sequences randomly selected from the entire study cohort. The threshold values were tuned using 25% of the annotation data. The detection accuracy (i.e., true positive, false positive, and false negative) was estimated with the rest of annotations which was not used for the parameter tuning (Fig. 3c). We obtained the Precision-Recall curve (Fig. 3d) and estimated the optimal F1 and Cohen’s kappa which were 0.80 and 0.77, respectively. These scores satisfied common criteria for the inter-observer reliability in behavioral sciences^34^, thus indicating that our automated analysis was reliable enough for quantification of social behavior. Furthermore, we used this detector for the rest of the entire dataset and found that the food-sharing event occurred more frequently in male than in female parents (Fig. 3e-f). Such a difference between fathers and mothers is suggested by previous studies on distinct species of New World monkeys^35–37^. The overall results demonstrated that our AI-based analytic pipeline clarified the differential roles of cooperating breeding animals in parenting under the laboratory environment, and that this pipeline could be useful for quantifying social behavior.

**Fig. 3.**
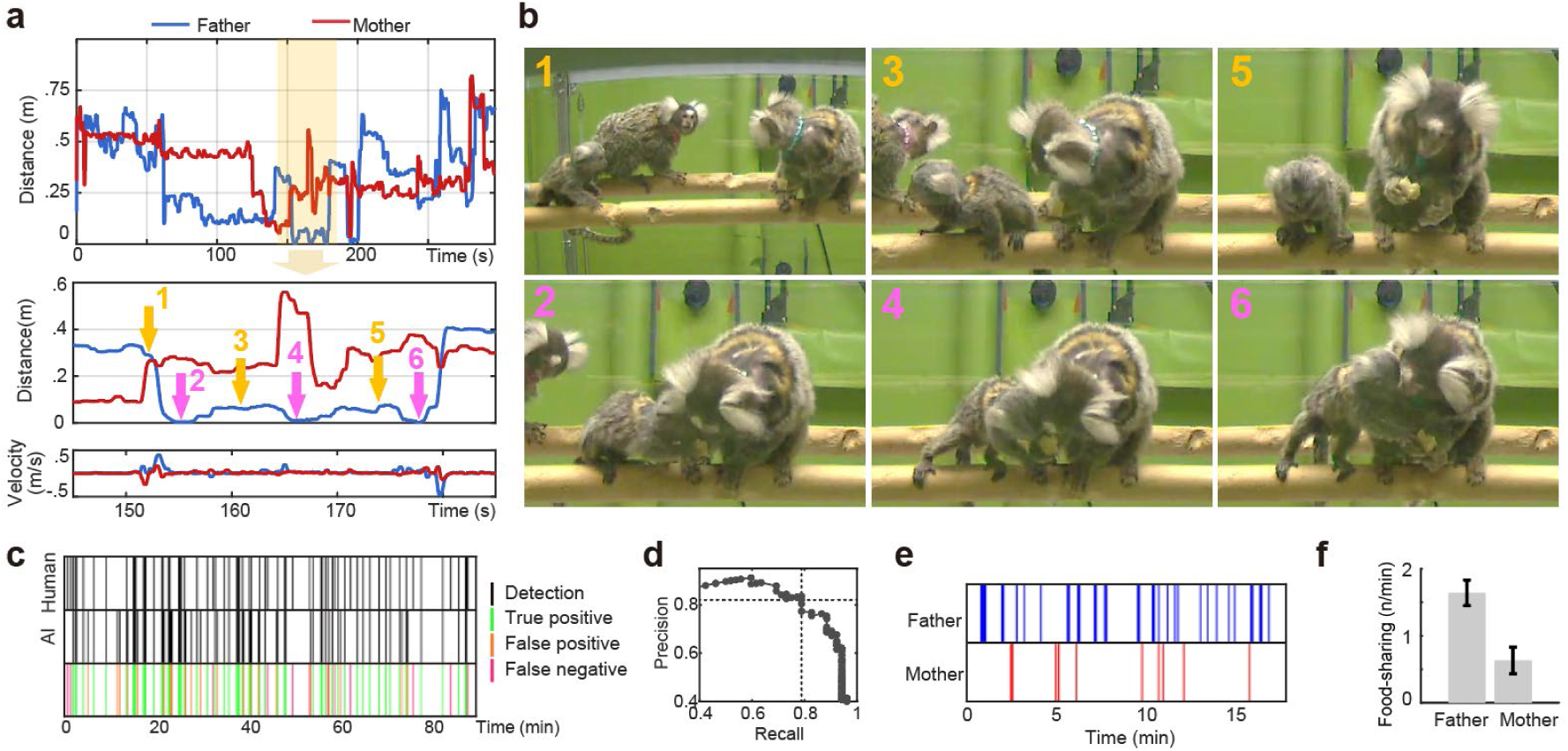
Differential contributions of father vs. mother marmosets to food-sharing events with their infants. **a:** Time-course of engineered features obtained from 3D poses of a marmoset family to predict food-sharing events. Top, the distance between the infant’s hand (either left or right) or mouth and those of the parents (for details, see the Methods section). Middle, the magnified view of the top panel. Each number corresponds to that of each image in **b**. Bottom, the velocity calculated as the first derivative of the middle panel. **b:** Exemplified scenes taken from marmoset family recordings. Each number corresponds to that of the time point in **a**; neutral in yellow (1,3,5) and food-sharing events in magenta (2, 4, 6). Note that the smallest values in the mid-panel of **a** correspond to the food-sharing events. **c:** Comparison between the human annotations (as ground truth) and the AI-based prediction obtained by applying a spatiotemporal filter to the engineered features shown in **a**. Colors in the third row represent true positive, false-positive, and false negative, respectively. The most of AI detection were true positive (i.e., the green bars were predominant). **d:** Precision-recall curve of the optimized detection. The highest F1 value (0.80) was at the intersection of the two dotted lines. This detection performance satisfied a common criterion in animal behavior research. **e:** Food-sharing events between the male/female and their infant predicted from the AI-based analysis for an example session. **f:** Rates of food-sharing events averaged across days. Our behavioral quantification via the AI-based pipeline reveals that the food- sharing event with infants occurs more frequently in male than in female parents (*df*=25, *t*=4.55, *p*=0.0001).

### Behavioral adjustment depending on others’ internal state as investigated by recurrent neural networks

In the second set of our experiments, we assessed the extent to which our system with 3D pose time-course data could infer the animal’s internal state behind actions. In social life of primates, it is crucial to adjust one’s own behavior depending on others’ internal state, such as emotions, intentions, and other physiological needs^38–40^. Conceivably, internally-guided behavioral changes by others may not readily be observable, but can be judged by watching over themselves^41^. Several human neuroimaging studies have shown neural substrates that are involved in this sort of cognitive function^42–45^. On the other hand, only a few related works have so far been available in NHPs^46–48^, because nonverbal behavioral paradigms are so limited that the possible underlying mechanism remains to be investigated. Here, we attempted to overcome this issue by combining a novel freely-moving behavioral task with our analytic pipeline using a deep neural network.

To examine a social behavioral action in response to others’ internal state, we developed a food competition task under freely moving conditions where two marmosets interacted to share or keep a valuable food (Fig. 4a,b). Two different pairs of marmosets participated in this experiment. The partner’s internal state (either full or hungry) was controlled without notifying the subject before the experiment started. Then, only the subject animal could obtain a large food that takes a couple of minutes to eat. The partner animal in the same cage may try to take away or beg for the food from the subject, and, therefore, the subject should pay attention to the partner’s action. Employing this behavioral task, we tested how the subject might adjust his/her behavior depending on the partner’s internal state.

**Fig. 4.**
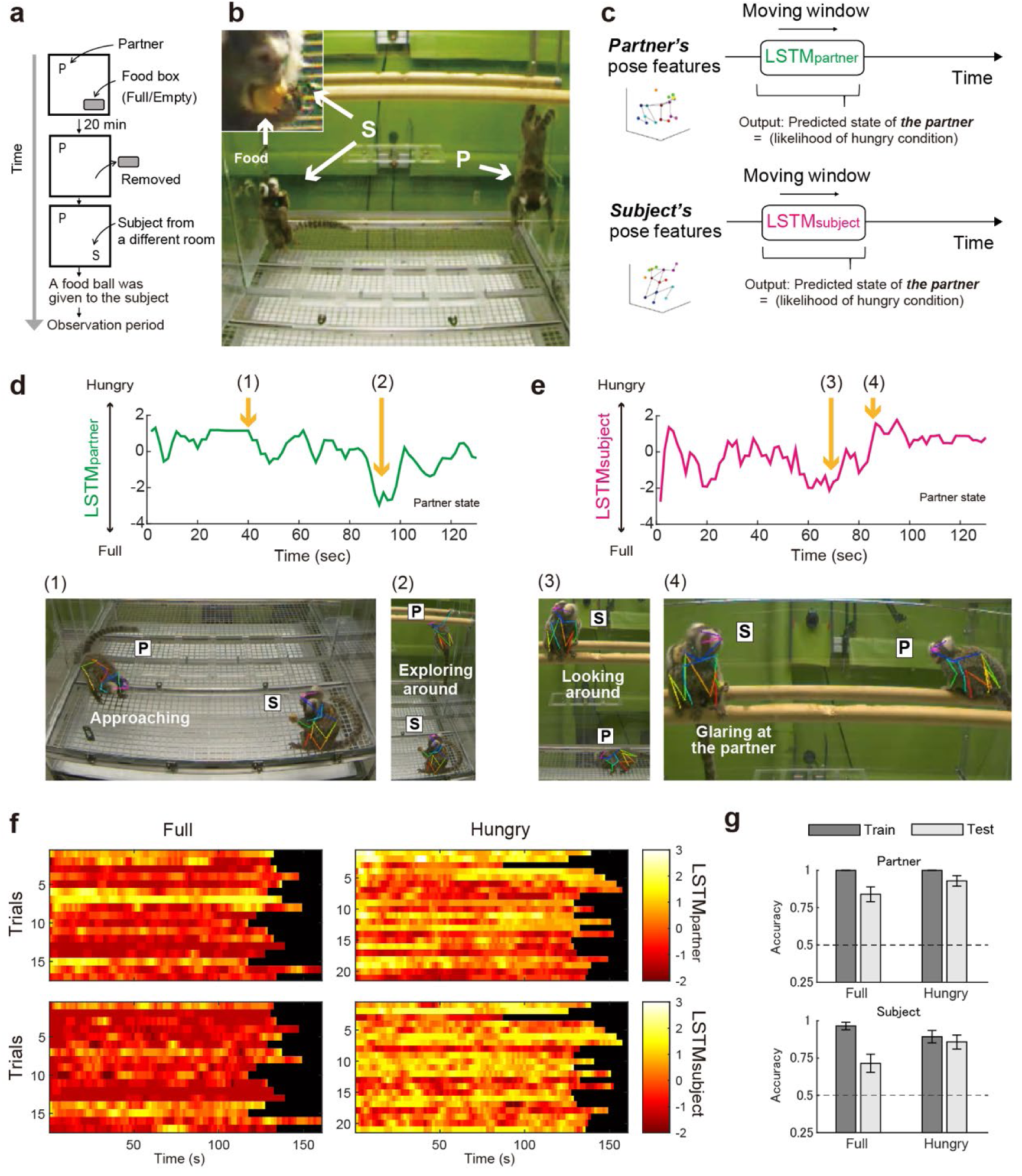
Quantification of the internal state predicted from actions. **a:** Design of a food-competition task. The partner (P)’s internal state (either full or hungry) was controlled before the main experiment without notifying the subject (S). **b:** Exemplified scene during the task and the magnified face of the subject (Inset). A valuable food (gum ball) was given only to the subject. The partner might simply display no interest in the food, or attempt to take away the food from the subject, and, therefore, the subject should pay attention to the partner. **c:** Schemes of analytic approaches using long-short-term-memory (LSTM) networks. Two different LSTMs with the same architecture, i.e., LSTM_partner_ and LSTM_subject_, were trained to predict the partner’s internal state from either the partner’s or the subject’s actions as input. **d and e:** Time-courses of the predicted internal state of the partner obtained as the output from the LSTM_partner_ and LSTM_subject_, respectively. Higher scores indicate the behavior in a hungrier state. Images in the lower panels are the scenes corresponding to the frames specified by the numbers in brackets in the upper row. Behavioral types indicated in white were determined by visual inspection of video recordings. Two examples shown here were derived from the hungry condition. **f:** Predicted internal state of the partner. Note that there are clear differences between the two conditions, while subtle variations within a trial or across days are also quantified. **g:** Performance of the LSTM_partner_ and LSTM_subject_ to discriminate conditions. The training and test of the networks were conducted in different subsets from the whole dataset. Shown is the average accuracy across all trials. Errors denote standard errors. The internal state of the partner can accurately be predicted not only from the actions of the partner itself, but also from those of the subject (binomial test; *ps*<0.0006, *n*=56 for each condition).

The Long Short-Term Memory (LSTM) ^49^, a type of recurrent neural network for temporal data analysis was used to decode the partner’s internal state. Two different LSTMs, LSTM_partner_ and LSTM_subject_, with the same architecture were trained to decode the partner’s internal state (i.e., full or hungry) from actions of either the subject or the partner (Fig. 4c). These LSTMs were designed to utilize the 3D pose data for 800ms as input and to generate as output a score representing the partner’s internal state, i.e., hungriness. The output score of LSTM_partner_ was predicted only from the partner’s action and could even display a variability within single trials (Fig. 4d). For example, in a scene with higher score (Fig. 4D, left panel), the partner was directly approaching the subject as if the partner tried to take away the food from the subject. Conversely, in a scene with lower score, the partner was exploring inside the cage without any interest in either the subject or the food. Similarly, as the partner’s internal state (and the resulting action) might probably affect the subject’s behavior, the output score of LSTM_subject_ was able to predict the partner’s internal state solely from the subject’s action (Fig. 4e). Even though the outputs of both LSTMs fluctuated within single trials or across trials, the overall scores were higher in a hungry than in a full condition (Fig. 4f). Thus, not only LSTM_partner_ but also LSTM_subject_ precisely predicted the partner’s condition on average (Fig. 4g). The accurate decoding of the LSTM_subject_ output indicated that the marmoset indeed adjusted his/her own behavior flexibly based on others’ internal state.

Another important question arises as to whether such a behavioral change might be an immediate, simple reaction to an others’ particular action rather than a reflection of others’ internal state behind the sequence of their actions. The comparison between the LSTM_partner_ and the LSTM_subject_ exhibited a positive correlation, which indicated that an immediate action by the subject was related to the sequence of the partner’s actions at that moment regardless of the partner’s internal state (Fig. 5a). Concurrently, at any level of the LSTM_partner_ output, the LSTM_subject_ output was consistently higher in the hungry than in the full condition (Fig. 5b). The present result implied that the one’s reaction towards the same sort of action by the other was changed according to the internal state. As an example of such behavioral adjustment depending on others’ internal state, we found that, in a pair of marmosets, the gaze behavior of the subject was changed according to the partner’s internal state. One marmoset sometimes looked back at the other when the other marmoset looked at the one (Fig. 5c). This look-back behavior was more frequently seen in a hungry than in a full condition (Fig. 5d), again indicating that the subject’s reaction towards the same action by the partner was changed based on the partner’s internal state. The overall results demonstrated the cognitive complexity of marmosets in the social context, thus elucidating that they flexibly adjust their behaviors depending on others’ internal state that is not readily observable by an immediate action alone.

**Fig. 5.**
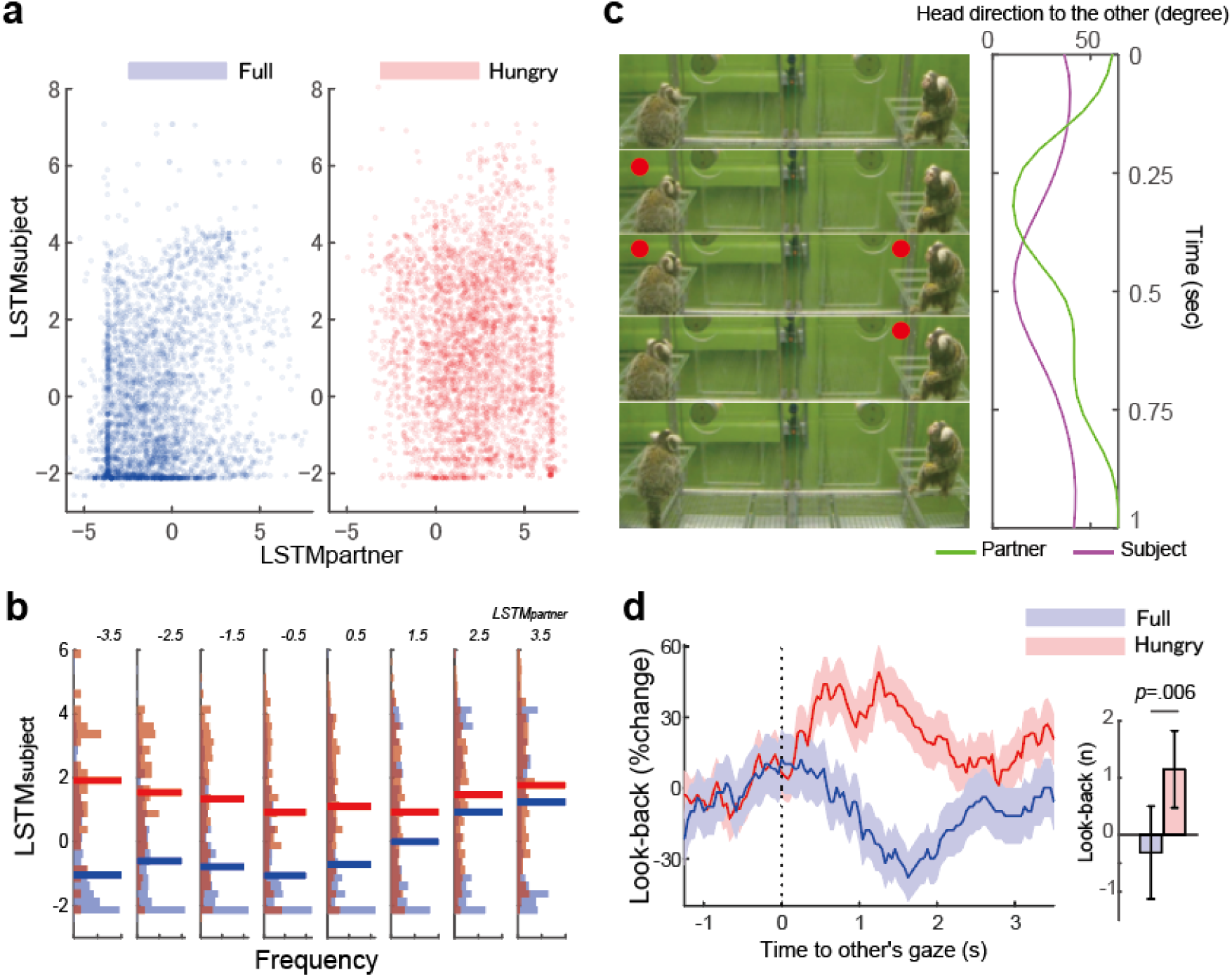
Response to partner’s action changes according to the internal state. **a:** Relationship between the instantaneous LSTM output of the partner and the subject. The likelihood of the partner’s hungry state predicted from the subject’s actions is positively correlated with that from the partner’s actions (*r*=0.22, *p*<0.001 and *r*=0.045, *p*=0.007 for the full and hungry condition respectively). Importantly, at the same level of the LSTM_partner_ output, the LSTM_subject_ output is substantially different across conditions. **b:** The partner’s internal state predicted from the subject’s actions at different levels of the LSTM_partner_ output. Light-colored bars represent the distributions of the LSTM_partner_ output, and thicken bars represent median. At all levels of the LSTM_partner_ output, the LSTM_subject_ output is significantly different according to the partner’s internal state (*ps* < 0.001) indicating that the subject’s response to similar actions by the partner changes according to the partner’s internal state.. **c:** Look-back behavior when two marmosets interact with each other. On the left side, five images are arranged in a time-series fashion from the top to the bottom. The left animal is the partner, and the right animal is the subject. Red circles indicate the timing when one marmoset was looking at the other. On the right side, a diagram shows time-dependent changes in the head angle of one marmoset towards the other. The partner directed the head towards the subject, and the subject looked back at the partner. **d:** Time-dependent changes in the subject’s gaze towards the partner aligned by the onset of the partner’s gaze. The y-axis represents the changes in the subject’s look-back behavior from baseline period (-1 ∼ 0s) . In a full (blue) or hungry (red) condition, the line and shaded zone represent the mean and standard error, respectively. The bar plots represent sum and standard error of look-back response between 0.5 to 1.5s, showing significant difference across conditions (*df*=555, *t*=2.77, *p*=0.0058) Note that the subject looked back at the partner more frequently in the hungry than in the full condition.

### Progressive manifestation of motor deficits in a marmoset model of PD as revealed by unsupervised clustering

In the third set of our experiments, we evaluated whether a completely unsupervised approach might allow us to detect behavioral changes in response to pathological conditions if relatively large-scale 3D pose data are available. To this end, we analyzed symptomatic behaviors in a marmoset model of PD. It is well known that PD progressively manifests motor deficits, such as akinesia, rigidity, and tremor, which is caused by degeneration/loss of dopaminergic neurons in the substantia nigra pars compacta (SNc) ^50,51^. Given that over-expression of mutant variants of alpha-synuclein (α-syn) emulates the progressive aspect of the disease, much emphasis has been placed on the notion that an animal model produced by α-syn over-expression is suitable for PD research^52,53^. In this model, however, observations over months or even years are required for behavioral assessment of phenotype expression, and, therefore, automated quantification of symptomatic behaviors is indispensable. Here, we yielded a PD model marmoset by injecting a combination of adeno-associated virus (AAV) vector^54^ carrying the mutant α-syn gene^55,56^ and pathological α-syn fibril^57^ into the nigra on one side of the brain (Fig. 6a). Histological analysis using tyrosine hydroxylase (TH) immunostaining after the behavioral observation confirmed loss of dopaminergic neurons from the SNc. With this PD model, varying motor activity was monitored for two days per month over one year.

**Fig. 6.**
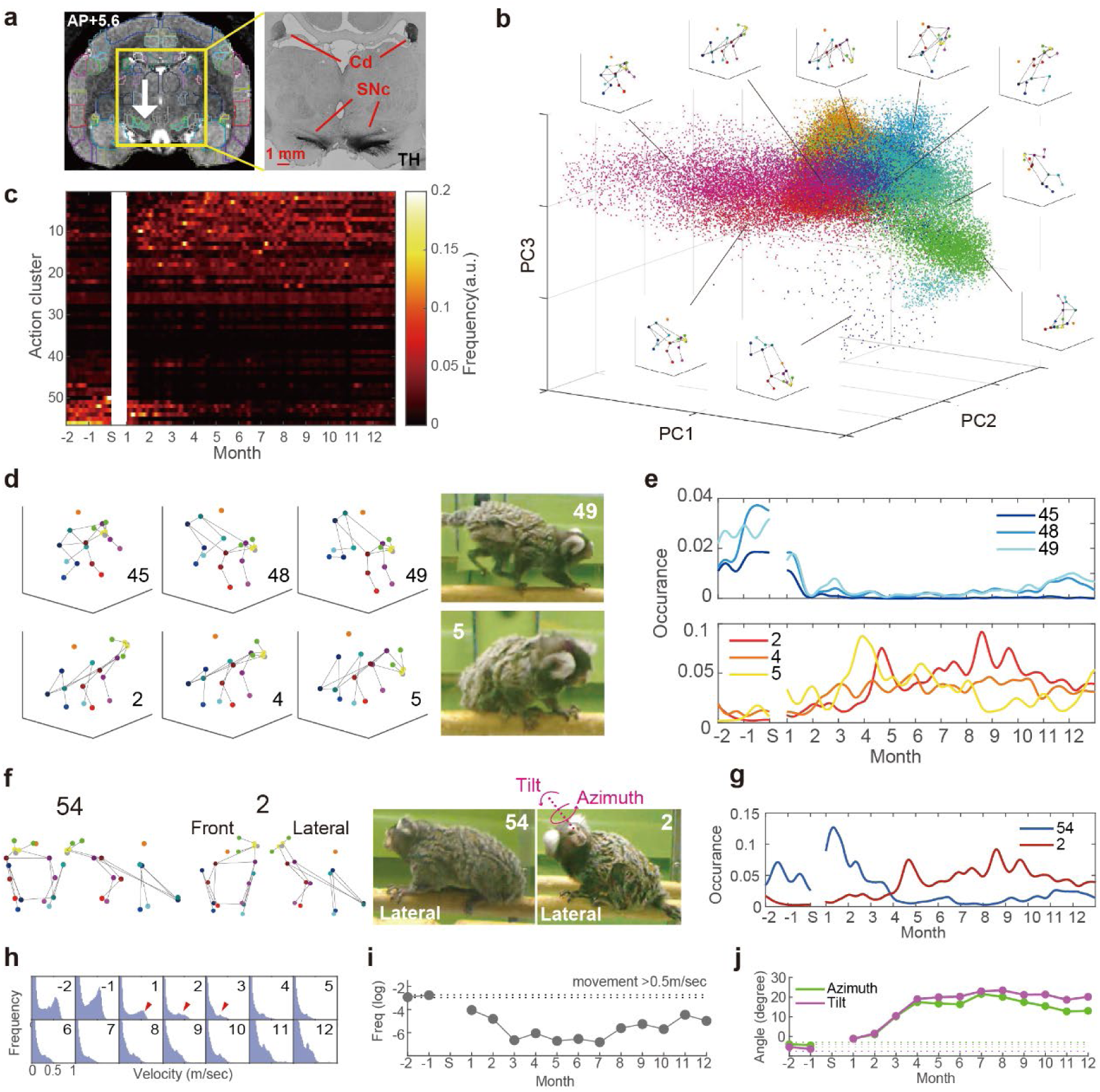
Longitudinal evaluation of progressive motor symptoms in a PD model marmoset. **a:** Left, site of combined injections of AAV vector carrying the mutant α-syn gene and pathological α-syn fibril seed in the substantia nigra compacta (SNc) indicated by the white arrow. Natural behaviors were recorded on two consecutive days per month before and after the injections. Right, section of tyrosine hydroxylase (TH) staining showing the lateralized degradation of TH in the SNc and the caudate, a projection target of the SNc. **b:** Results of the unsupervised action motif clustering. Each dot represents an instance of action which was plotted on the axes of the first three principal components (PC1-3) used for the action classification, and each color corresponds to the classified action. **c:** Heatmap representing the frequency of each action motif cluster observed over longitudinal recordings from the pre-injections to 12 months post-injections. **d:** Examples of the most frequently observed actions before and after the injections. Before the injections (cluster 45, 48 and 49), quick locomotion, such as jumping, gallop, and quick turning, was mainly observed. In contrast, a variety of “stay” postures with slight differences were primarily seen after the injections (2, 4 and 5). Exemplified images were taken from the videographic data. Each number corresponds to an action cluster in **c**. **e:** Time-course for the frequency of occurrence of action clusters shown in **d**. **f:** Representative postures which seem similar though clustered into different classes. Note the differences in the tilt angle and azimuth of the head between the pre-injection (i.e., 54) and the post-injection (i.e., 2). Front, frontal view. Lateral, lateral view. **g:** Time-course for the frequency of occurrence of action clusters shown in **f**. **h:** Histograms of the locomotion speed in different months. Red triangles represent a decreasing trend of fast locomotion as the progression of motor symptoms. **i:** Analysis of bradykinesia as assessed with changes in the frequency of fast locomotion (0.5 m/s) over several months. The dotted horizontal lines indicate upper and lower bound of the 95% confidential interval for pre-injections data (note that these bounds were so close so that it is hard to see the gap). **j:** Analysis of abnormal posture in the neck as assessed with changes in the tilt angle and azimuth of the head over several months. The dotted horizontal lines indicate 95% confidential as in **I** (again, the bounds were so close).

Employing our analytic pipeline in a fully unsupervised manner without any a priori hypothesis, we could identify a couple of behavioral changes in the marmoset PD model. First, by means of dimensional reduction and clustering approach, we determined action motifs that were the patterns of 3D pose time-series data repeatedly observed throughout the recording period (Fig. 6b; for details, see the Methods section). We found that some actions were occasionally observed before the surgery, and others gradually appeared after the surgery (Fig. 6c). Specific behavioral actions, such as running, turning, and jumping from wood, were reduced after the surgery (Fig. 6d,e; upper panels). Conversely, various types of “stay” actions were increasingly observed several months after the surgery (Fig. 6d,e; bottom panels). Interestingly, apparently similar postures were classified into different clusters notably by the difference in the neck angle (Fig. 6f). Some postures were seen more frequently, whereas others were observed less frequently after the surgery (Fig, 6g). After three months, an increased tonus of the neck muscle markedly appeared contralaterally as evidenced by the finding that the head bent towards the side opposite to the nigral injection site.

We further quantified the amount of gross movement (as an index of reduced locomotion) and the head posture based on the 3D pose time-series data, and then successfully confirmed the progression of symptomatic behaviors obtained from the unsupervised analysis (Fig. 6h-j). The overall results indicated that parkinsonian phenotypes induced by α-syn over-expression gradually progressed. This suggested that our system allowed the longitudinal and high-throughput evaluation of symptomatic behaviors in brain disease models without any behavioral tasks.

## Discussion

In the present study, we have developed the analytic pipeline that permits automated and high-throughput quantification of natural behaviors of common marmosets using a markerless motion capture system which consists of multiple deep neural networks. With the large-scale ground truth dataset, the decoding accuracy reached the best performance that we could expect at the present time. Applying this system, we have revealed that our approach is capable of detecting behavioral changes due to a variety of experimental conditions, such as differential contributions of males vs. females to parenting in breeding families, flexible behavioral adjustment depending on others’ internal state, and progressive manifestation of motor impairments in a PD model. Our results provide a novel framework to many research areas of neuroscience using NHPs by introducing objective and large-scale quantification of animal behavior. It should also be noted here, however, that there are some limitations on the use of the analytic pipeline that we have developed in this study. First, the proposed system is able to quantify only restricted variations of behavioral actions that are represented by 18 keypoints. Thus, other types of actions, such as facial expression, cannot be quantified^62^. Second, careful assessment is needed to confirm that behavioral data obtained from our system are not attributable to erroneous tracking of individual animals. The 3D pose time-course data may sometimes be derived from a mixture of multiple animals, although such an error is rare as shown in Figure 2c,d. In a severe condition where individual recognition is inaccurate, an alternative system should be called for to address this issue specifically^58^.

Recent technological innovations have attracted much attention to experimental paradigms with freely moving marmosets. Large-scale telemetric recordings of neuronal activity were successfully carried out^31^, and electrocorticography recordings from almost the entire lateral hemisphere were also reported^59,60^. Combining these recording techniques with our analytic pipeline allows comprehensive understanding of the correlation between cortical signals and behavioral dynamics. This could be an appropriate methodology to explore the cortical circuitry related to behavioral actions of particular interest. Then, optogenetic^61^ and chemogenetic^62^ approaches, which are also compatible with freely-moving experimental conditions, enable us to disclose the causal role of a specific neural circuit in the expression of a given type of natural behaviors. Until recently, major efforts have been made to assess motor and cognitive functions of NHPs through analysis of eye/hand movements as the behavioral output. Now, the AI-based innovative development has increasingly been accomplished to quantify and evaluate social interactions in a certain animal population with high efficiency^3^. This may make it feasible to elucidate the neural mechanisms underlying behavioral theories, so far intensively explored in socio-ecological and ethological studies, for example, the Machiavellian theory in which expansion of the cerebral cortex, especially the frontal lobe, leads to the adaptation to social complexity in our daily life^63–65^.

The novel pipeline that we have established for 3D pose time-series analysis of a group of marmosets can be utilized in various experimental environments and laboratories. All that is required is to estimate the camera calibration parameters for accurate 3D reconstructions and to refine the neural networks for detection, identification and pose estimation of individuals. Concerning the former requirement, at least a two-camera system should work though our experiments were conducted with eight cameras to enhance the robustness and accuracy, and then data needed for the calibration will be acquired within hours. With respect to the latter requirement, the neural networks for 2D analysis should be re-tuned to each experimental environment or laboratory because of the differences in varying factors, such as background, lighting, and camera angle. In our experiments, we provided a substantial amount of ground truth data to achieve robust 3D analysis, which will be of immense help for adapting neural networks to specific environments and achieving impeccable performance. In recent years, several tools, for example, “style transfer” ^66,67^, further support a transfer learning of the networks from some environment to others. Moreover, while our analytic pipeline has highly been optimized for marmosets, It can be customized for other species as well.

The present study has revealed the potent applicability of the 3D pose data, as evidenced by a wide range of behavioral parameters beyond spatiotemporal kinematics that can be quantified via a proper choice of downstream analytic methodologies (Fig. 7). The simplest method is to detect specific behavioral events by defining spatiotemporal parameters derived from certain combinations of 3D keypoints, as demonstrated in the food-sharing experiment. A key factor to succeed in this method is appropriate feature engineering that is suitable for target event detection and parameter tuning with a small set of supervised data, both of which should be performed by experts of animal behavioral observations. Moreover, we have elucidated that simple spatiotemporal data concerning the 3D poses permit quantification of the internal state of marmosets which is combined with cutting-edge neural networks, for instance, a recurrent neural network (i.e., LSTM) in the present study. This brings about a unique opportunity of studying the mind behind the complex social behavior in primates. Finally, a fully-unsupervised data mining approach is capable of disclosing behavioral changes induced by pathophysiological manipulation, as shown in the PD model experiment. This approach is specifically beneficial to explore behavioral changes comprehensively if a substantial amount of data are available. Such methodological innovations are greatly meritorious given that the behavioral complexity inherent in NHPs substantially accentuates the assessment of neurological/psychiatric/developmental disorder models. The high-throughput and versatile trait of our evaluation system will play critical roles in establishing a new standard that quantifies social/pathophysiological behaviors of NHPs.

**Fig. 7.**
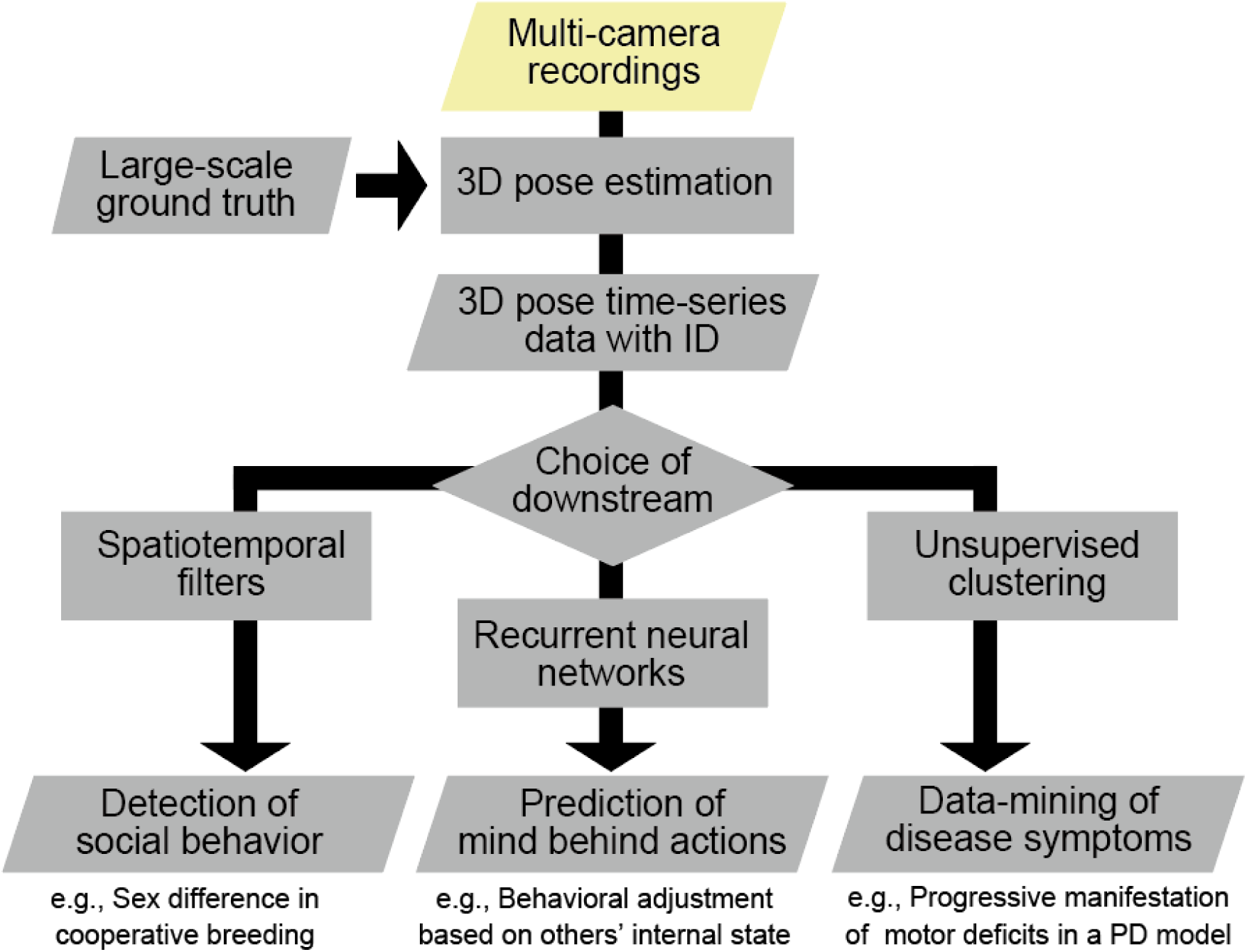
Overview of comprehensive approaches to quantification of marmoset natural behaviors based on 3D poses. Analytic workflow in the present study. The 3D pose time-course data *per se* are merely pieces of spatiotemporal information about body postures involving the limb positions. However, by combining with proper downstream analytic methodologies, the data allow us to elucidate a wide spectrum of behavioral parameters based on the 3D poses alone.

## Methods

### Animals

All procedures for the use and experiments of common marmosets were approved by the Animal Welfare and Animal Care Committee of the Center for the Evolutionally Origins of the Human Behavior, Kyoto University, followed by the Guidelines for Care and Use of Nonhuman Primates established by the same institution. First, 29 marmosets (ranging from 1.5 months olds to 12 years old; 13 males and 16 females) were used to create the ground truth dataset. Four adult and two infant marmosets derived from two families participated in the food-sharing experiment. Then, two pairs of adult marmosets were utilized for the food competition experiment, and one adult marmoset was for the PD model experiment.

### Recording system

A recording booth was a 90-cm cubic box which consisted of acrylic transparent walls, and a metal mesh floor and ceiling. This recoding booth was designed to keep up to four animals under the Ethical Guideline of the Japan Neuroscience Society and equipped with common items required for a normal marmoset cage, such as water bottles, food boxes, and perches. Videographic images were recorded by Motif system (Loopbio, Lange G, Wien, Austria) which was synchronized with eight machine vision cameras (2048x1536-pixel, 24 fps). The cameras were arranged horizontally with an equal distance as surrounding the recording booth. The viewing angle of each camera was set at 110x70 degree to cover the whole booth.

To accomplish accurate 3D reconstructions, we obtained intrinsic (e.g., lens distortion coefficients) and extrinsic (e.g., camera positions) camera calibration parameters by the OpenCV framework as follows: The intrinsic parameters were obtained by cv2.omnidir.calibrate using the images of a checker-board pattern recorded by each camera; and the extrinsic parameters were initialized by cv2.solvPnP function by the 3D coordinates of a set of landmark positions in the recording booth and their 2D coordinates projected onto the camera image. To improve the calibration accuracy, we further optimized both the intrinsic and the extrinsic parameters simultaneously by minimizing the projection (reconstruction) errors of the trajectory of a small object (a ping-pong ball) moved inside the recording cage^68^.

### Ground truth dataset

Our keypoint schema follows that of macaque-pose^12^ dataset with slight modification to fit to analyze the whole-body movements of marmosets. Specifically, we annotated 20 keypoints consisting of the nose, eyes (left and right), ears, shoulders, elbows, wrists, hips, knees, ankles, back, and the middle and tip of the tail (while the last two keypoints were not used in the analytic pipeline). The annotators were trained by movies of marmosets whose body parts corresponding to the keypoints were marked by paint markers. The annotations were performed in a 3D manner by using custom-made software where those of a single body were a collection of 3D positions constructed through triangulation of 2D positions via all cameras. While the 3D positions could be computed with triangulation once a single keypoint was annotated via more than two cameras, the annotators visually confirmed every keypoint for all cameras to maximize precision. We used images from 29 different marmosets and annotated 7404 bodies in a 3D space which were equivalent to 56103 bodies in a 2D space. We selected scenes from different behavioral contexts, 732 bodies from full-day recordings of a single animal, 654 bodies from those of two animals, 2010 bodies from three-animal recordings, and 4008 bodies from four-animal recordings. The annotation frames were semi-manually selected to maximize variations of the behavioral contents.

### Markerless 3D multi-animal pose estimation

The analytic pipeline started from the analysis of 2D images taken from each camera (Fig.1b *2D processes*). The detection network analyzed the locations of marmosets in an image of each frame and generated a bunch of bounding boxes, which are rectangles of partial regions bounded by the smallest rectangle enclosing a marmoset as a region of interest. Then, the pose network estimated 18 keypoints, and the ID network estimated an animal ID for all bounding boxes. The bounding boxes were combined along the time axis at the 2D level to construct so-called 2D tracklets, namely time-series data consisting of the regions of a marmoset associated with the postures and animal IDs. As multiple bounding boxes could be detected in each frame, the bounding boxes that seemed to correspond to a single marmoset were combined based on the consistency in the positions of the marmoset across frames. At this moment, the 2D tracklets were still fragmented in short durations, because one animal who were occluded by objects or other animals, and, therefore, it could not be tracked continuously. The 2D processing was implemented using OpenMMLab^69^, a set of image processing libraries for deep neural networks. The network architecture used here was yolox-l^70^ and resnet-50^71^ for detection and identification. The pose networks were hrnet-w32^72^ for both the food-sharing and the food competition experiments, and dekr-hrnet-w48^73^ for the PD model experiment. The connections of bounding boxes to construct 2D tracklets were performed by using *ByteTrack*^74^.

Subsequently, the 3D pose time-series data on each animal were obtained with four steps. The first to third steps corresponded to *camera association* and the fourth step to *3D optimization* in Figure 1a.

As the first step, in each frame, we grouped the bounding boxes (tracklets were not used here) likely belonging to the same marmosets across different cameras. This process was performed only in key frames which were every 0.5 sec to reduce computational load. We searched for the optimal grouping of bounding boxes by minimizing geometric inconsistency (i.e., the inverse of the so-called pose affinity score^75^) between the boxes from different cameras within a group. We defined geometric inconsistency *D_g_* as below.

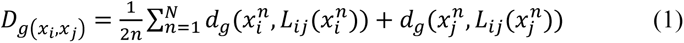

where *x_i_^n^* indicated the 2D position of the *n*-th keypoint of pose *I*, *L_ij_*(*x_j_^n^*) the projection line associated with *x_j_^n^* from a different camera, and *d*_*g*_(⋅, *l*) the point-to-line distance for l. The optimization was performed according to the algorithm proposed by Dong et al. ^80^. Once the grouping of bounding boxes was established, we constructed the 3D pose of a marmoset for each group of the bounding boxes by triangulation of the 2D poses in each key frame. Then, we obtained 3D poses of marmosets in every key frame, while their temporal association remained undetermined.

As the second step, the matching of the same animal over time was performed as follows: A combination of 3D poses across adjacent key frames could be considered, in the Graph theory, the maximum matching *M* of a complete bipartite graph *G=(S,T;E)* with non-negative edge cost *c*: *E* → ℝ ≥ 0, where *S*, *T* are 3D poses for key frame *t* and *t+*1. Here we defined the cost *c(i,j)* for the edge connecting *Si* and *Tj* poses as below.

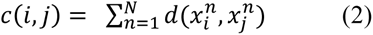

where *N* indicated the number of keypoints, *x_i_^n^* and *x_j_^n^* represented the 3D position of the *n*-th keypoint of the 3D pose *Si* and *Tj*, respectively, and *d*(⋅,⋅) was the distance between two points in a 3D space. This cost represented geometrical inconsistency of a pair of 3D poses.

The maximum matching *M* was obtained by minimizing the cost ∑_*e*∈*M*_ *c*(*e*) through the Hungarian algorithm. In addition, the edge connections were removed if the geometrical inconsistency per keypoint was over an empirically determined threshold *T_1_*=150. The frames between the key frames were complemented by continuity of 2D tracklets which were combinations of multiple bounding boxes over time in a 2D space. Through this process, we obtained 3D tracklets time-series data on 3D posture.

Third, a marmoset ID was assigned for each 3D tracklet. The ID was assigned in every frame if the following criterion was satisfied:

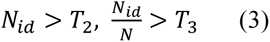

where *N_id_* was the number of instances for the most frequently observed ID, *N* was the number of all bounding boxes taken from all cameras, *T_2_* and *T_3_* were hyperparameters, set as 12 and 0.8, respectively. Here, within a sliding time window (5 sec), all the bounding boxes belonging to a single 3D tracklet were considered. If the same ID was assigned to a different tracklet at the same time point, the ID was given only to the tracklet with the highest *N_id_*. A 3D tracklet was divided at the time point when the IDs assigned by the above criterion were changed within the 3D tracklet.

The fourth step was the final refinement of the 3D tracklets. There might be the case where multiple 3D tracklets, which should correspond to the same animal, were dissociated due to the failure in the previous steps. To compensate such a case, these tracklets were integrated by the following procedure. Suppose that there was a tracklet that had not yet been assigned an ID, *T_noID_*; and a tracklet that had been assigned an ID, *T_withID_*. During the period when two 3D tracklets overlapped, if the difference between their 3D trajectory was less than the error threshold *T*_4_=200, then the ID of *T_withID_* propagated to that of *T_noID_*. This was repeated twice for the entire dataset. Furthermore, for tracklets that had not yet been assigned an ID, we assigned the remaining ID if the IDs of all but one animal had been assigned. Finally, the tracklets with the same ID were integrated, and the resulting 3D pose time-series data on individual marmosets were spatiotemporally smoothed and normalized via anipose^73^.

### Food-sharing experiment

A couple of marmoset families participated in this experiment. Each family consisted of a father, a mother, and their infant who was about three months of age at the start of the experiment. A piece of home-made Arabian gumball was given to each of the parents simultaneously, and then their social interactions were observed. When both gumballs were consumed, new ones were given again to the parents separately. The experiment was carried out for about 30 min per day and repeated for 12 or 16 days in two families.

Food-sharing events were detected by the following procedure. First, 3D pose data on three individuals per family were obtained with the analytic pipeline. At this stage, the 3D pose data were independent across the animals and were not suitable for detecting social behavior. Therefore, we created new features by combining the 3D pose data about the infant and parents. Specifically, we calculated the distance between the mouth or the left or right hand of the infant and those of each parent. Considering all combinations for a pair of the infant and one of his/her parent, this process generated 9 different values for each time frame. The smallest one of these values was taken for each frame, and the resulting time-series data (*D*) and the first derivative (*V*) were obtained. A food-sharing event was marked when there were at least *T_N_* consecutive frames in which *D* and *V* were larger than detection parameters *T_d_* and *T_v_*. To optimize these detection parameters and to evaluate detection accuracy, a human observer counted the occurrence of food-sharing events as a subset of the entire dataset. The human observer coded the presence or absence of food-sharing events for every 15 sec and analyzed for 90 min in total. The threshold value obtained from the human coding was optimized by maximizing the consistency to the automated detection by using 25% of the annotation data. The detection accuracy was obtained from the rest of the annotation data. The Precision-Recall curve shown in Figure 3d was obtained by varying *T_d_* from the optimal value. The statistical significance in the difference between the father and the mother in the food-sharing events were evaluated with a paired two-tailed *t*-test (α = 0.05) with the number of observations on each recording day as independent data points.

### Food competition experiment

Two pairs of marmosets were used for this experiment. For each pair, the subject and partner animals were familiar with each other as they had been kept in the same cage. Food deprivation was performed from the evening of one day before the experiment, and, therefore, both animals were in a hungry state at the start of the experiment. Just before the experiment, the partner’s state was controlled to be either hungry or full by the following procedure. The partner was separated from the subject immediately before the experiment in order that they could not see each other. In a full condition, enough food was provided until the partner could not eat any more. In a hungry condition, the partner was forced to stay for the same duration as in the full condition. In each trial, the subject but not the partner was provided with a gumball with high reward value, and their social interactions were observed. The experiments were performed 3- 4 trials per day, and a hungry or a full condition was randomly assigned across days.

LSTM^49^, a type of recurrent neural network, was used to predict the partner’s internal state obtained from 3D pose data on either the subject or the partner. We coded LSTMs using the implementation in pytorch 2.0. The architecture of LSTMs for the subject and partner was the same. The LSTMs took 3D pose time-course data for 20 frames (corresponding to 800 ms) from either the partner or the subject as input and generated two output scores indicating that the likelihood of the partner’s internal state was either hungry or full. Cross entropy loss was computed across the outputs and the experimental conditions for the network training. As a quantitative representation of the partner’s hungriness (such as in Fig.4d), we took the value in the final full connection layer before the softmax function.

The input dataset for LSTM networks was composed of the aligned-posture, locomotion speed, degree of approach-avoidance, and head direction, as calculated by the following procedure. To obtain the aligned-posture, the 3D pose data were shifted frame by frame, and, thus, the midpoint between the left and the right hip keypoint was aligned in the same position across the frames. Then, the aligned data were further rotated along the horizontal plane, and, therefore, the azimuth of the trunk was aligned across the frames. The locomotion speed was the first derivative of the trajectory of the hip-mid point. The approach-avoidance was the inner product of the locomotion vector (a vector connecting the mid-point of the hip keypoint across adjacent frames) and the vector from the position of one marmoset to that of another marmoset. The head direction was the angle between the one’s head direction (the 45-degree upright vector from a vector connecting the nose and the midpoint of the left and right eyes and ears) and the direction to the other.

One fourth of the total data was used to train the networks, and the training was iterated until the learning curve reached a plateau. The best network weights over the training iterations were selected based on the performance of the prediction for the rest of the dataset which had not been used for the training. In the analysis of Figure 5c and d, the look-back behavior was defined as the head direction (as defined in the previous paragraph) of the subject (as the calculation mentioned above) became below 40 degrees and were aligned by the onset of the partner’s gaze (head direction should be below 40 degrees) which was kept for more than 800 ms. The bar graph in Figure 5d denoted the sum of the look-back behavior between 0.5-1.5 sec to the partner’s gaze. The statistical significance was obtained by an unpaired two-sided *t*-test for the differences between the conditions.

### PD model experiment

A PD marmoset model was produced by unilateral injections of both virus vector expressing mutant α-syn and pathological α-syn fibril into the SNc. A total of 12-μl solutions consisting of 4 μl of AAV2.1-hTH-α-syn (G51D) (4.88x10e13 gc/ml) ^54^ and 8 μl of the fibril (5 mg/ml) ^57^ was injected into four rostrocaudally and mediolaterally different loci of the SNc through a 10-μl Hamilton microsyringe (30 gauge) over 35 min per penetration. The injection coordinates were adjusted individually based on MR images. A surgical navigation system (Brainsight, Rogue Research, Montréal, Québec, Canada) was used to accurately guide the position of the injection sites^76^. The animal was anesthetized with ketamine hydrochloride (20- 40 mg/kg, i.m.) and maintained with isoflurane (1-2%) during the surgery while SpO_2_, heart rate, and rectal temperature were monitored. A water-heating circulator was used to control the body temperature. An analgesic (Meloxicam; 0.1-0.2 mg/kg, i.m.) was also administered before and for a couple of days after the injection. Behavioral observations were conducted once a month. The marmoset was moved to the recording booth and allowed to stay there for two days. Food pellets were supplied once a day and water was available *ad libitum*. Video recordings was done for 20 min per hour from 9 a.m. to 4 p.m. (a total of 160 min per day). The recordings were started two months before the surgery and continued 12 months after the surgery.

After the behavioral assessment, immunohistochemical analysis was performed to confirm loss of dopamine neurons from the SNc. The animal was deeply anesthetized with ketamine hydrochloride (40mg/kg, i.m.) and sodium secobarbital (50 mg/kg, i.v.), and perfused transcardially with 0.1M phosphate-buffered saline (PBS) followed by 4% paraformaldehyde in 0.1 M phosphate buffer (pH 7.4). Then, the brain was removed from the skull, postfixed overnight, and saturated with 30% sucrose at 4°C. Coronal sections were cut serially at the 40- µm thickness on a freezing microtome. A series of every tenth section was used for tyrosine hydroxylase (TH) immunostaining. The sections were pretreated with 0.3% H_2_O_2_ for 30 min and immersed in 1% skim milk for 2 hr. The sections were then incubated for 48 hr at 4°C with mouse anti-TH antibody (1:2,000; Millipore, Burlington, MA) in 0.1 M PBS containing 2% normal donkey serum and 0.1% Triton X-100. Subsequently, the sections were incubated with biotinylated donkey anti-mouse IgG antibody (1:1,000; Jackson ImmunoResearch, West Grove, PA) for 2 hr at room temperature in the same fresh medium, followed by the avidin-biotin-peroxidase complex (ABC Elite; 1:200; Vector laboratories, Burlingame, USA) in 0.1 M PBS for 2 hr at room temperature. Finally, the antigen was visualized with diaminobenzidine (DAB) containing nickel ammonium sulfate (0.01% DAB, 1.0% nickel ammonium sulfate, and 0.0003% H_2_0_2_). The sections were mounted onto gelatin-coated glass slides and counterstained with 1% Neutral red.

An unsupervised clustering of behavioral actions was performed by using time-series data about action features which were computed based on the 3D pose data as follows: First, the aligned postures were obtained as described in the previous section. Then, the spectrogram representation (0.05-12.8 Hz) of these data was obtained from the fast Fourier transformation, and, therefore, the data at a single time point contained not only instantaneous postural information, but also dynamics of the postures. In the end, the action features used for the clustering were created by adding locomotion vector to this spectrogram. The clustering was carried out by using the k-means clustering method with the number of classes fixed ====to 56, and, thus, all videographic frames throughout the entire recording period were classified as one of the 56 action clusters. Then, the time-course of the occurrence rate of each action class was obtained as shown in Figure 6c. The order of these action clusters was defined by the following procedure. The 56-dimensional time-series data representing the action occurrence rate were analyzed by the principal component analysis (PCA). Then, the first principal component *PC1* showed monotonic increment in which the score was low before the surgery and was gradually being increased after the surgery. Therefore, the order of the coefficients of *PC1* was used as the order of the action clusters. In other words, the actions with the small cluster number were frequently observed after the surgery, and those with the larger cluster number were often observed before the surgery. In Figure 6j, the azimuth and tilt of the head were calculated by the vector form the midpoint of the shoulders to that of the eyes in the aligned posture. For both the movements and the head angles, the errors were estimated by the bootstrap method. All data during the pre-surgery period were used to estimate the 95% confidential intervals. The mean for every 15 min was taken as an independent data point, and the repetition of the bootstrap sampling was 2000 times.

## Acknowledgements

We are grateful to Eri Sumiya for the care of animals, Akihisa Kaneko and Dr. Takako Miyabe for their veterinary support, Maki Fujiwara and Mayuko Nakano for preparation of the virus vector, Dr. Cesar Aguirre for preparation of the fibril, and Emiko Tanaka for technical support of the histological analysis. We also thank Amarbayasgalant Badarch for annotations of animal identification and tracking. This work was supported by JSPS KAKENHI Grant Numbers 19H05467 to M.T., 22H05157 and 22K19480 to Ki.I., and 22K07325 to T.K.; AMED Brain/MINDS Grant Number JP22dm0207077 to M.T. and T.K.

## Author contributions

T.K., J.M., Ki.I. and M.T. designed the experiments. T.K. and J.M. developed the analytic pipeline. T.K., W.L., X.Z., L.U., K.K., Y.O. performed the experiments. A.Z. and Ki.I. prepared the viral vector. Ks.I., K.B. and H.M. prepared the fibril. T.K. J.M. and Ki.I. analyzed data. H.N., T.O., Ki.I. and M.T. supervised the experimenters. T.K. and M.T. wrote the draft. T.K., J.M., Ki.I. and M.T. reviewed and edited the manuscript.

## Declaration of interests

The authors declare no competing interests.

## Data and code availability

The 3D ground truth for marmosets and code for the core functions will be made available in a public repository at the time of publication in a peer reviewed journal. Additional code and data associated with this work is available from the Lead Contact upon request.

## Notes

### Competing Interest Statement

The authors have declared no competing interest.

## References

1. Coffey, K.R., Marx, R.E., and Neumaier, J.F. (2019). DeepSqueak: a deep learning-based system for detection and analysis of ultrasonic vocalizations. Neuropsychopharmacology 44, 859–868.

2. Dolensek, N., Gehrlach, D.A., Klein, A.S., and Gogolla, N. (2020). Facial expressions of emotion states and their neuronal correlates in mice. Science 368, 89–94.

3. Schofield, D., Nagrani, A., Zisserman, A., Hayashi, M., Matsuzawa, T., Biro, D., and Carvalho, S. (2019). Chimpanzee face recognition from videos in the wild using deep learning. Sci. Adv. 5, eaaw0736.

4. Wu, Z., Zhang, C., Gu, X., Duporge, I., Hughey, L.F., Stabach, J.A., Skidmore, A.K., Hopcraft, J.G.C., Lee, S.J., Atkinson, P.M., et al. (2023). Deep learning enables satellite-based monitoring of large populations of terrestrial mammals across heterogeneous landscape. Nat. Commun. 14, 3072.

5. Graving, J.M., Chae, D., Naik, H., Li, L., Koger, B., Costelloe, B.R., and Couzin, I.D. (2019). DeepPoseKit, a software toolkit for fast and robust animal pose estimation using deep learning. eLife 8, e47994.

6. Lauer, J., Zhou, M., Ye, S., Menegas, W., Schneider, S., Nath, T., Rahman, M.M., Di Santo, V., Soberanes, D., Feng, G., et al. (2022). Multi-animal pose estimation, identification and tracking with DeepLabCut. Nat. Methods 19, 496–504.

7. Mathis, A., Mamidanna, P., Cury, K.M., Abe, T., Murthy, V.N., Mathis, M.W., and Bethge, M. (2018). DeepLabCut: markerless pose estimation of user-defined body parts with deep learning. Nat. Neurosci. 21, 1281–1289.

8. Pereira, T.D., Tabris, N., Matsliah, A., Turner, D.M., Li, J., Ravindranath, S., Papadoyannis, E.S., Normand, E., Deutsch, D.S., Wang, Z.Y., et al. (2022). SLEAP: A deep learning system for multi-animal pose tracking. Nat. Methods 19, 486–495.

9. Marshall, J.D., Aldarondo, D.E., Dunn, T.W., Wang, W.L., Berman, G.J., and Ölveczky, B.P. (2021). Continuous Whole-Body 3D Kinematic Recordings across the Rodent Behavioral Repertoire. Neuron 109, 420–437.e428.

10. Schneider, A., Zimmermann, C., Alyahyay, M., Steenbergen, F., Brox, T., and Diester, I. (2022). 3D pose estimation enables virtual head fixation in freely moving rats. Neuron 110, 2080–2093.e2010.

11. Berger, M., Agha, N.S., and Gail, A. (2020). Wireless recording from unrestrained monkeys reveals motor goal encoding beyond immediate reach in frontoparietal cortex. eLife 9, e51322.

12. Voloh, B., Maisson, D.J.N., Cervera, R.L., Conover, I., Zambre, M., Hayden, B., and Zimmermann, J. (2023). Hierarchical action encoding in prefrontal cortex of freely moving macaques. Cell Rep. 42, 113091.

13. Ebina, T., Obara, K., Watakabe, A., Masamizu, Y., Terada, S.-I., Matoba, R., Takaji, M., Hatanaka, N., Nambu, A., Mizukami, H., et al. (2019). Arm movements induced by noninvasive optogenetic stimulation of the motor cortex in the common marmoset. Proc. Natl. Acad. Sci. U. S. A. 116, 22844–22850.

14. Labuguen, R., Matsumoto, J., Negrete, S.B., Nishimaru, H., Nishijo, H., Takada, M., Go, Y., Inoue, K.-i., and Shibata, T. (2021). MacaquePose: A Novel “In the Wild” Macaque Monkey Pose Dataset for Markerless Motion Capture. Front. Behav. Neurosci. 14, 581154.

15. Shaw, L., Wang, K.H., and Mitchell, J. (2023). Fast prediction in marmoset reach-to-grasp movements for dynamic prey. Curr. Biol. 33, 2557–2565.e2554.

16. Yao, Y., Bala, P., Mohan, A., Bliss-Moreau, E., Coleman, K., Freeman, S.M., Machado, C.J., Raper, J., Zimmermann, J., Hayden, B.Y., and Park, H.S. (2023). OpenMonkeyChallenge: Dataset and Benchmark Challenges for Pose Estimation of Non-human Primates. Int. J. Comput. Vis. 131, 243–258.

17. Testard, C., Tremblay, S., Parodi, F., DiTullio, R.W., Acevedo-Ithier, A., Gardiner, K., Kording, K.P., and Platt, M. (2023). Neural signatures of natural behavior in socializing macaques. bioRxiv, 2023.2007.2005.547833.

18. Miller, C.T., Freiwald, W.A., Leopold, D.A., Mitchell, J.F., Silva, A.C., and Wang, X. (2016). Marmosets: A Neuroscientific Model of Human Social Behavior. Neuron 90, 219–233.

19. Yano-Nashimoto, S., Truzzi, A., Shinozuka, K., Murayama, A., Kurachi, T., Moriya-Ito, K., Tokuno, H., Miyazawwa, E., Esposito, G., Okano, H., et al. (2023). Infant attachment behaviors reflect the parenting style of individual caregiver in common marmosets. bioRxiv, 2023.2005.2018.541258.

20. Huang, J., Cheng, X., Zhang, S., Chang, L., Li, X., Liang, Z., and Gong, N. (2020). Having Infants in the Family Group Promotes Altruistic Behavior of Marmoset Monkeys. Curr. Biol. 30, 4047–4055.e4043.

21. Courellis, H.S., Nummela, S.U., Metke, M., Diehl, G.W., Bussell, R., Cauwenberghs, G., and Miller, C.T. (2019). Spatial encoding in primate hippocampus during free navigation. PLoS Biol. 17, e3000546.

22. Samandra, R., Haque, Z.Z., Rosa, M.G.P., and Mansouri, F.A. (2022). The marmoset as a model for investigating the neural basis of social cognition in health and disease. Neurosci. Biobehav. Rev. 138, 104692.

23. Okano, H. (2021). Current Status of and Perspectives on the Application of Marmosets in Neurobiology. Annu. Rev. Neurosci. 44, 27–48.

24. Kishi, N., Sato, K., Sasaki, E., and Okano, H. (2014). Common marmoset as a new model animal for neuroscience research and genome editing technology. Dev. Growth Differ. 56, 53–62.

25. Feng, G., Jensen, F.E., Greely, H.T., Okano, H., Treue, S., Roberts, A.C., Fox, J.G., Caddick, S., Poo, M.M., Newsome, W.T., and Morrison, J.H. (2020). Opportunities and limitations of genetically modified nonhuman primate models for neuroscience research. Proc. Natl. Acad. Sci. U. S. A. 117, 24022–24031.

26. Sasaki, E., Suemizu, H., Shimada, A., Hanazawa, K., Oiwa, R., Kamioka, M., Tomioka, I., Sotomaru, Y., Hirakawa, R., Eto, T., et al. (2009). Generation of transgenic non-human primates with germline transmission. Nature 459, 523–527.

27. Sato, K., Oiwa, R., Kumita, W., Henry, R., Sakuma, T., Ito, R., Nozu, R., Inoue, T., Katano, I., Sato, K., et al. (2016). Generation of a Nonhuman Primate Model of Severe Combined Immunodeficiency Using Highly Efficient Genome Editing. Cell Stem Cell 19, 127–138.

28. Walker, J.D., Pirschel, F., Sundiang, M., Niekrasz, M., MacLean, J.N., and Hatsopoulos, N.G. (2021). Chronic wireless neural population recordings with common marmosets. Cell Rep. 36, 109379.

29. Roy, S., and Wang, X. (2012). Wireless multi-channel single unit recording in freely moving and vocalizing primates. J. Neurosci. Methods 203, 28–40.

30. Hoffmann, K., Coolen, A., Schlumbohm, C., Meerlo, P., and Fuchs, E. (2012). Remote long-term registrations of sleep-wake rhythms, core body temperature and activity in marmoset monkeys. Behav. Brain Res. 235, 113–123.

31. Bala, P.C., Eisenreich, B.R., Yoo, S.B.M., Hayden, B.Y., Park, H.S., and Zimmermann, J. (2020). Automated markerless pose estimation in freely moving macaques with OpenMonkeyStudio. Nat. Commun. 11, 4560.

32. Wang, J., Tan, S., Zhen, X., Xu, S., Zheng, F., He, Z., and Shao, L. (2021). Deep 3D human pose estimation: A review. Comput. Vis. Image Underst. 210, 103225.

33. Brown, G.R., Almond, R.E.A., and Bergen, Y.v. (2004). Begging, stealing, and fffering: food transfer in nonhuman primates. Adv. Stud. Behav. 34, 265–295.

34. Joseph, F.L., Bruce, L., and Paik, M.C. (2003). Statistical methods for rates and proportions (John Wiley & Sons, Inc.).

35. Saito, A., Izumi, A., and Nakamura, K. (2008). Food transfer in common marmosets: parents change their tolerance depending on the age of offspring. Am. J. Primatol. 70, 999–1002.

36. Guerreiro Martins, E.M., Moura, A.C.A., Finkenwirth, C., Griesser, M., and Burkart, J.M. (2019). Food sharing patterns in three species of callitrichid monkeys (Callithrix jacchus, Leontopithecus chrysomelas, Saguinus midas): Individual and species differences. J. Comp. Psychol. 133, 474–487.

37. Price, E.C., and Feistner, A.T.C. (2001). Food Sharing in Pied Bare-Faced Tamarins (Saguinus bicolor bicolor): Development and Individual Differences. Int. J. Primatol. 22, 231–241.

38. Call, J., and Tomasello, M. (2008). Does the chimpanzee have a theory of mind? 30 years later. Trends Cogn. Sci. 12, 187–192.

39. Isoda, M. (2021). The Role of the Medial Prefrontal Cortex in Moderating Neural Representations of Self and Other in Primates. Annu. Rev. Neurosci. 44, 295–313.

40. Isoda, M., Noritake, A., and Ninomiya, T. (2018). Development of social systems neuroscience using macaques. Proc. Jpn. Acad. Ser. B Phys. Biol. Sci. 94, 305–323.

41. Lokesh, R., Sullivan, S., Calalo, J.A., Roth, A., Swanik, B., Carter, M.J., and Cashaback, J.G.A. (2022). Humans utilize sensory evidence of others’ intended action to make online decisions. Sci. Rep. 12, 8806.

42. Tso, I.F., Rutherford, S., Fang, Y., Angstadt, M., and Taylor, S.F. (2018). The “social brain” is highly sensitive to the mere presence of social information: An automated meta-analysis and an independent study. PLoS One 13, e0196503.

43. Frith, C.D., and Frith, U. (2006). The neural basis of mentalizing. Neuron 50, 531–534.

44. Gallagher, H.L., and Frith, C.D. (2003). Functional imaging of ‘theory of mind’. Trends Cogn. Sci. 7, 77–83.

45. Siegal, M., and Varley, R. (2002). Neural systems involved in ’theory of mind’. Nat. Rev. Neurosci. 3, 463–471.

46. Hayashi, T., Akikawa, R., Kawasaki, K., Egawa, J., Minamimoto, T., Kobayashi, K., Kato, S., Hori, Y., Nagai, Y., Iijima, A., et al. (2020). Macaques Exhibit Implicit Gaze Bias Anticipating Others’ False-Belief-Driven Actions via Medial Prefrontal Cortexortex. Cell Rep. 30, 4433–4444.e4435.

47. Haroush, K., and Williams, Ziv M. (2015). Neuronal prediction of opponent’s behavior during cooperative social interchange in primates. Cell 160, 1233–1245.

48. Roumazeilles, L., Schurz, M., Lojkiewiez, M., Verhagen, L., Schüffelgen, U., Marche, K., Mahmoodi, A., Emberton, A., Simpson, K., Joly, O., et al. (2021). Social prediction modulates activity of macaque superior temporal cortex. Sci. Adv. 7, eabh2392.

49. Hochreiter, S., and Schmidhuber, J. (1997). Long Short-Term Memory. Neural Comput. 9, 1735–1780.

50. Obeso, J.A., Rodriguez-Oroz, M.C., Rodriguez, M., Lanciego, J.L., Artieda, J., Gonzalo, N., and Olanow, C.W. (2000). Pathophysiology of the basal ganglia in Parkinson’s disease. Trends Neurosci. 23, S8–S19.

51. Chiken, S., Takada, M., and Nambu, A. (2021). Altered Dynamic Information Flow through the Cortico-Basal Ganglia Pathways Mediates Parkinson’s Disease Symptoms. Cereb. Cortex 31, 5363–5380.

52. Shimozawa, A., Ono, M., Takahara, D., Tarutani, A., Imura, S., Masuda-Suzukake, M., Higuchi, M., Yanai, K., Hisanaga, S.-i., and Hasegawa, M. (2017). Propagation of pathological α-synuclein in marmoset brain. Acta Neuropathol. Commun. 5, 12.

53. Eslamboli, A., Romero-Ramos, M., Burger, C., Bjorklund, T., Muzyczka, N., Mandel, R.J., Baker, H., Ridley, R.M., and Kirik, D. (2007). Long-term consequences of human alpha-synuclein overexpression in the primate ventral midbrain. Brain 130, 799–815.

54. Kimura, K., Nagai, Y., Hatanaka, G., Fang, Y., Tanabe, S., Zheng, A., Fujiwara, M., Nakano, M., Hori, Y., Takeuchi, R.F., et al. (2023). A mosaic adeno-associated virus vector as a versatile tool that exhibits high levels of transgene expression and neuron specificity in primate brain. Nat. Commun. 14, 4762.

55. Lesage, S., Anheim, M., Letournel, F., Bousset, L., Honoré, A., Rozas, N., Pieri, L., Madiona, K., Dürr, A., Melki, R., et al. (2013). G51D α-synuclein mutation causes a novel Parkinsonian–pyramidal syndrome. Ann. Neurol. 73, 459–471.

56. Kiely, A.P., Asi, Y.T., Kara, E., Limousin, P., Ling, H., Lewis, P., Proukakis, C., Quinn, N., Lees, A.J., Hardy, J., et al. (2013). α-Synucleinopathy associated with G51D SNCA mutation: a link between Parkinson’s disease and multiple system atrophy? Acta Neuropathol. 125, 753–769.

57. Hayakawa, H., Nakatani, R., Ikenaka, K., Aguirre, C., Choong, C.-J., Tsuda, H., Nagano, S., Koike, M., Ikeuchi, T., Hasegawa, M., et al. (2020). Structurally Distinct α-Synuclein Fibrils Induce Robust Parkinsonian Pathology. Mov. Disord. 35, 256–267.

58. Matsumoto, J., Kaneko, T., Kimura, K., Negrete, S.B., Guo, J., Suda-Hashimoto, N., Kaneko, A., Morimoto, M., Nishimaru, H., Setogawa, T., et al. (2023). Three- dimensional markerless motion capture of multiple freely behaving monkeys for automated characterization of social behavior. bioRxiv, 2023.2009.2013.556332.

59. Kaneko, T., Komatsu, M., Yamamori, T., Ichinohe, N., and Okano, H. (2022). Cortical neural dynamics unveil the rhythm of natural visual behavior in marmosets. Commun. Biol. 5, 108.

60. Komatsu, M., Kaneko, T., Okano, H., and Ichinohe, N. (2019). Chronic Implantation of Whole-cortical Electrocorticographic Array in the Common Marmoset. Journal of Visualized Experiments 144, e58980.

61. Yoshimoto, S., Araki, T., Uemura, T., Nezu, T., Sekitani, T., Suzuki, T., Yoshida, F., and Hirata, M. (2016). Implantable wireless 64-channel system with flexible ECoG electrode and optogenetics probe. 2016 IEEE Biomedical Circuits and Systems Conference (BioCAS), 476–479.

62. Mimura, K., Nagai, Y., Inoue, K.-i., Matsumoto, J., Hori, Y., Sato, C., Kimura, K., Okauchi, T., Hirabayashi, T., Nishijo, H., et al. (2021). Chemogenetic activation of nigrostriatal dopamine neurons in freely moving common marmosets. iScience 24, 103066.

63. Dunbar, R.I.M. (1992). Neocortex size as a constraint on group-size in primates. J. Hum. Evol. 22, 469–493.

64. Dunbar, R.I.M., and Shultz, S. (2021). Social complexity and the fractal structure of group size in primate social evolution. Biol. Rev. 96, 1889–1906.

65. Whiten, A., and Byrne, R.W. (1988). Tactical deception in primates. Behav. Brain Sci. 11, 233–244.

66. Bolaños, L.A., Xiao, D., Ford, N.L., LeDue, J.M., Gupta, P.K., Doebeli, C., Hu, H., Rhodin, H., and Murphy, T.H. (2021). A three-dimensional virtual mouse generates synthetic training data for behavioral analysis. Nat. Methods 18, 378–381.

67. Kim, J., Kim, M., Kang, H., and Lee, K. (2019). U-GAT-IT: Unsupervised Generative Attentional Networks with Adaptive Layer-Instance Normalization for Image-to-Image Translation. arXiv, arXiv:1907.10830.

68. Karashchuk, P., Rupp, K.L., Dickinson, E.S., Walling-Bell, S., Sanders, E., Azim, E., Brunton, B.W., and Tuthill, J.C. (2021). Anipose: A toolkit for robust markerless 3D pose estimation. Cell Rep. 36, 109730.

69. Chen, K., Wang, J., Pang, J., Cao, Y., Xiong, Y., Li, X., Sun, S., Feng, W., Liu, Z., Xu, J., et al. (2019). MMDetection: Open MMLab Detection Toolbox and Benchmark. arXiv:1906.07155.

70. Ge, Z., Liu, S., Wang, F., Li, Z., and Sun, J. (2021). YOLOX: Exceeding YOLO Series in 2021. arXiv:2107.08430.

71. He, K., Zhang, X., Ren, S., and Sun, J. (2015). Deep Residual Learning for Image Recognition. arXiv:1512.03385.

72. Sun, K., Xiao, B., Liu, D., and Wang, J. (2019). Deep High-resolution Representation Learning for Human Pose Estimation. arXiv:1902.09212.

73. Geng, Z., Sun, K., Xiao, B., Zhang, Z., and Wang, J. (2021). Bottom-Up Human Pose Estimation Via Disentangled Keypoint Regression. arXiv:2104.02300.

74. Zhang, Y., Sun, P., Jiang, Y., Yu, D., Weng, F., Yuan, Z., Luo, P., Liu, W., and Wang, X. (2021). ByteTrack: Multi-Object Tracking by Associating Every Detection Box. arXiv:2110.06864.

75. Dong, J., Jiang, W., Huang, Q., Bao, H., and Zhou, X. (2019). Fast and Robust Multi- Person 3D Pose Estimation from Multiple Views. arXiv:1901.04111.

76. Mundinano, I.-C., Flecknell, P.A., and Bourne, J.A. (2016). MRI-guided stereotaxic brain surgery in the infant and adult common marmoset. Nat. Protoc. 11, 1299–1308.

